# Genotype-dependent and non-gradient patterns of RSV gene expression

**DOI:** 10.1101/641878

**Authors:** Felipe-Andrés Piedra, Xueting Qiu, Michael N. Teng, Vasanthi Avadhanula, Annette A. Machado, Do-Kyun Kim, James Hixson, Justin Bahl, Pedro A. Piedra

## Abstract

Respiratory syncytial virus (RSV) is a nonsegmented negative-strand (NNS) RNA virus and a leading cause of severe lower respiratory tract illness in infants and the elderly. Transcription of the ten RSV genes proceeds sequentially from the 3’ promoter and requires conserved gene start (GS) and gene end (GE) signals. Previous studies using the prototypical GA1 genotype Long and A2 strains have indicated a gradient of gene transcription. However, recent reports show data that appear inconsistent with a gradient. To better understand RSV transcriptional regulation, mRNA abundances from five RSV genes were measured by quantitative real-time PCR (qPCR) in three cell lines and cotton rats infected with virus isolates belonging to four different genotypes (GA1, ON, GB1, BA). Relative mRNA levels reached steady-state between four and 24 hours post-infection. Steady-state patterns were genotype-specific and non-gradient, where mRNA levels from the G (attachment) gene exceeded those from the more promoter-proximal N (nucleocapsid) gene across isolates. Transcript stabilities could not account for the non-gradient patterns observed, indicating that relative mRNA levels more strongly reflect transcription than decay. While the GS signal sequences were highly conserved, their alignment with N protein in the helical ribonucleocapsid, i.e., N-phase, was variable, suggesting polymerase recognition of GS signal conformation affects transcription initiation. The effect of GS N-phase on transcription efficiency was tested using dicistronic minigenomes. Ratios of minigenome gene expression showed a switch-like dependence on N-phase with a period of seven nucleotides. Our results indicate that RSV gene expression is in part sculpted by polymerases that initiate transcription with a probability dependent on GS signal N-phase.

**Author Summary:** RSV is a major viral pathogen that causes significant morbidity and mortality, especially in young children. Shortly after RSV enters a host cell, transcription from its nonsegmented negative-strand (NNS) RNA genome starts at the 3’ promoter and proceeds sequentially. Transcriptional attenuation is thought to occur at each gene junction, resulting in a gradient of gene expression. However, recent studies showing non-gradient levels of RSV mRNA suggest that transcriptional regulation may have additional mechanisms. We show using RSV isolates belonging to four different genotypes that gene expression is genotype-dependent and one gene (the G or attachment gene) is consistently more highly expressed than an upstream neighbor. We hypothesize that variable alignment of highly conserved gene start (GS) signals with nucleoprotein (i.e., variable GS N-phase) can affect transcription and give rise to non-gradient patterns of gene expression. We show using dicistronic RSV minigenomes wherein the reporter genes differ only in the N-phase of one GS signal that GS N-phase affects gene expression. Our results suggest the existence of a novel mechanism of transcriptional regulation that might play a role in other NNS RNA viruses.

## Introduction

Respiratory syncytial virus (RSV) can infect individuals repeatedly and is the most common pathogen associated with severe lower respiratory tract disease in children worldwide [1–5]. Numerous host-related and environmental risk factors for severe disease are known [6–8] while viral factors are less clear.

RSV is a nonsegmented negative-strand (NNS) RNA virus classified into two major subgroups, A and B, largely distinguished by antigenic differences in the attachment or G protein [9, 10]. The two subgroups are estimated to have diverged from an ancestral strain over 300 years ago [11] and have evolved into multiple co-circulating genotypes [11–15].

The RNA genome of RSV is embedded in interlinking and helix-forming subunits of nucleocapsid (N) protein, together forming the ribonucleocapsid (RNP) complex [16, 17]. Viral mRNA are not encapsidated [17, 18]. Formation of the RNP complex requires high concentrations of N protein and a 5’ terminal dinucleotide AC synthesized by the polymerase independently of template [18, 19]. Each subunit of N protein binds a seven nucleotide stretch of RNA via contacts with the sugar phosphate backbone, causing the RNA to adopt a conformation with a distinct configuration of solvent-exposed and buried nucleobases [16, 17]. Exposed nucleobases can presumably interact directly with viral polymerases bound to the RNP complex [16]. Moreover, the alignment of the viral RNA to the N protein within the RNP (N-phase) will determine its pattern of exposed and buried bases. The effects of N-phase on promoter recognition have been explored in RSV and some paramyxoviruses [18, 20–23]. N-phase affects RNA synthesis by paramyxoviral RNA polymerases but, in RSV, promoter recognition is strongly determined by the proximity of the promoter sequence to the 3’ terminus of the genome; replication is abolished if the core promoter starts six or more nucleotides from the 3’ end [23]. Unlike its effects on promoter recognition and replication, the effects of N-phase on transcription are unexplored.

Transcription in RSV and other NNS viruses is sequential, with genes transcribed in their order of occurrence from the 3’ promoter of the genome [18, 24–29]. Each of the ten genes of RSV contains essential gene start (GS) and gene end (GE) signals flanking the open reading frame (ORF) [30–32]. Transcription is initiated at the GS signal which also serves as a capping signal on the 5’ end of the nascent mRNA [18, 33, 34]. The polymerase then enters elongation mode until it reaches a GE signal, where the mRNA is polyadenylated and released [18, 30]. Two genes overlap at the 5’ end of the RSV genome. The GE signal of matrix 2 (M2) occurs downstream of the GS signal of the large polymerase (L) gene. The polymerase must return from the M2 GE signal for full-length L mRNA to be made [35], suggesting that transcribing polymerases scan the RSV genome bidirectionally for a new GS signal after terminating transcription. Indeed, scanning polymerase dynamics may be a universal feature of NNS virus transcription [18, 36–38].

By homology with other NNS viruses, it is widely assumed that transcription in RSV follows a gradient, where the extent to which a gene is transcribed falls with its distance from the 3’ promoter [29, 39, 40]. Earlier studies reported data consistent with a gradient [39, 41, 42]; however, recent studies show mRNA abundances that peak at the G gene, which is located in the middle of the genome [40, 43]. We recently reported the G gene to be the most abundant in clinical samples obtained from RSV/A- and RSV/B-infected infants [44]. Thus, existing data suggest that patterns of RSV gene expression are more variable than has been assumed.

Here we explored the natural diversity of patterns of RSV gene expression by using qPCR to measure mRNA abundances of five different RSV genes (NS1, NS2, N, G, F) from isolates that we sequenced belonging to both subgroups and four different genotypes (RSV/A/GA1, RSV/A/ON, RSV/B/GB1, RSV/B/BA). Genotype-dependent patterns were observed, all diverging from a gradient and all showing higher levels of G mRNA than expected. Transcript stabilities did not account for the non-gradient patterns of mRNA levels. We analyzed GS signal sequence and N-phase, and hypothesized that GS signal N-phase can affect RSV gene expression on the basis of our findings. We found evidence supporting this hypothesis by measuring gene expression from RSV minigenomes encoding two reporter genes, where minigenomes differed only in the N-phase of one GS signal.

## Results

### RSV mRNA abundances

Oligonucleotide standards of known concentration were used to convert cycle threshold (*C*_T_) values measured by real-time PCR for mRNA targets (Fig 1A) to mRNA abundances. Twenty oligonucleotide standards and sets of reagents (primers and probe) were needed to quantify 20 mRNA targets (five genes in four isolates). All reagents gave rise to a similar range of *C*_T_ values for standards at equal concentrations (Fig 1B).

**Fig 1.**
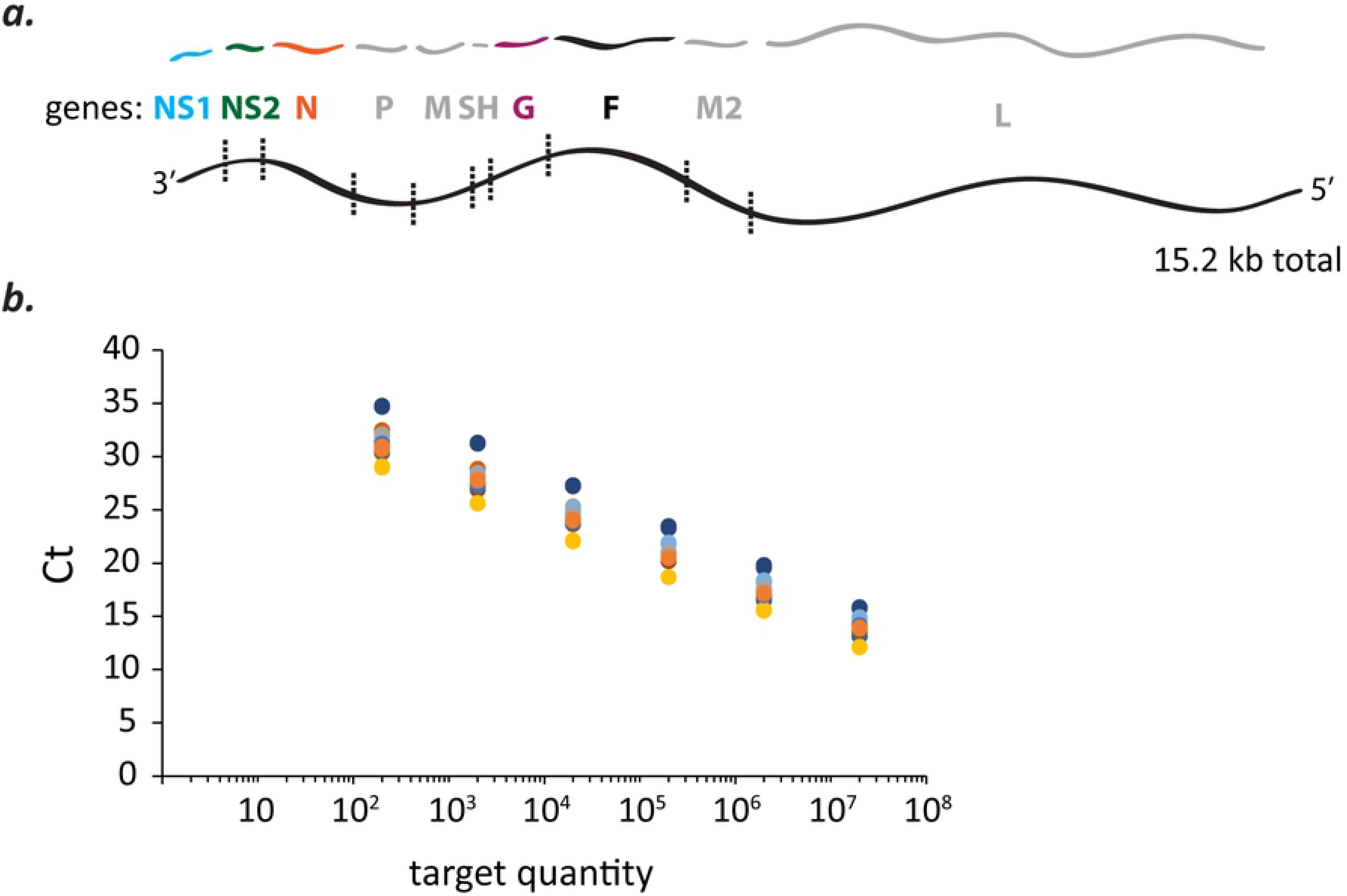
qPCR-based measurements of mRNA abundances from five RSV genes and four different virus isolates using oligonucleotide standards. **(a)** Five of 10 RSV genes were chosen for mRNA abundance measurements by qPCR. The 5 genes (NS1, NS2, N, G, & F) span half of the nucleotide length of the 15.2 kb genome and its entire gene length minus the final two genes, M2 and L. **(b)** Known amounts of different oligonucleotide standards were detected over a similar range of cycle threshold (*C*_T_) values. Twenty different oligonucleotide standards at known concentrations were needed (4 virus isolates x 5 mRNA targets) to convert *C*_T_ values measured for viral mRNA targets into mRNA abundances. Each dot represents the mean *C*_T_ of duplicate measurements of an oligonucleotide standard at a known concentration or quantity (= number of molecules / qPCR rxn). Dots of like color are dilutions of the same oligonucleotide standard.

### Relative mRNA levels

RSV isolates from both major subgroups (A and B) and four different genotypes (A/GA1/Tracy, A/ON/121301043A, B/GB1/18537, B/BA/80171) were used to infect HEp-2 cells (MOI = 0.01). Total mRNA abundances began to plateau at ∼48 hours post-infection (pi) for all isolates (Fig 2A), consistent with the presence of significant viral cytopathic effect beyond this time-point. Relative mRNA levels were calculated for each isolate at each time-point by dividing the abundance of each mRNA by the total mRNA abundance (Fig 2B). Relative mRNA levels reached steady-state between four and 24 hours pi (Fig 2B).

**Fig 2.**
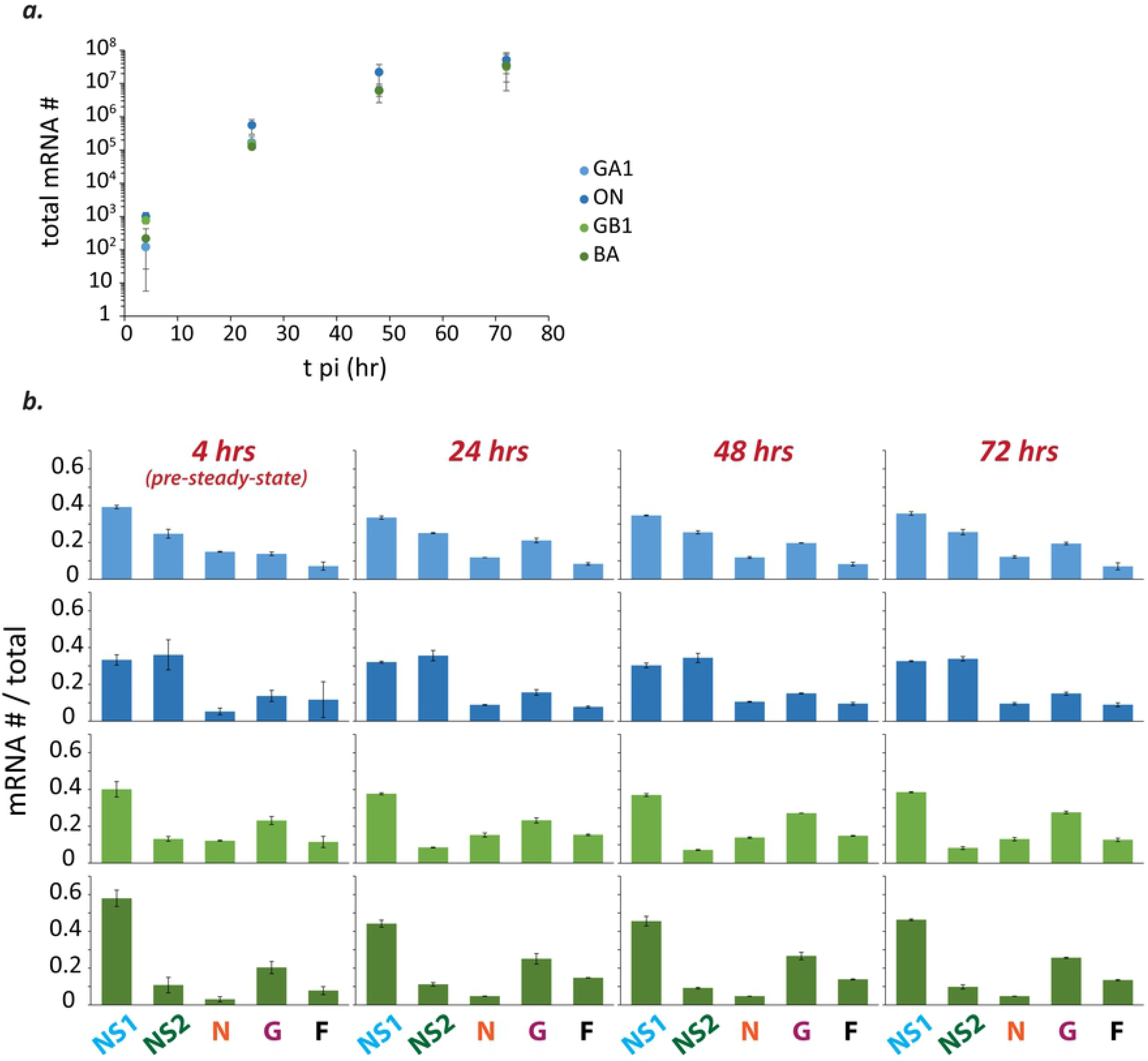
Total mRNA abundances plateau beyond 48 hours post-infection and relative mRNA levels reach steady-state soon after the start of infection. **(a)** Total mRNA abundances (= NS1+NS2+N+G+F) from HEp-2 cells infected with different isolates of RSV (MOI = 0.01) begin to plateau by ∼48 hours post-infection (pi). Each dot (RSV/A/GA1Tracy [pale blue]; RSV/A/ON/121301043A [dark blue]; RSV/B/GB1/18537 [light green]; RSV/B/BA/80171 [dark green]) represents the mean and error bars the standard deviation of two independent experiments (n=2). For each independent experiment, the mean was calculated from duplicate measurements and used in subsequent calculations. **(b)** Relative mRNA levels reach steady-state sometime between four and 24 hours pi. Histograms depicting relative mRNA levels are shown for all measured time-points (4, 24, 48, 72 hr pi) and all four isolates (color scheme same as (a)). Each bar depicts the mean mRNA # / total mRNA # of the indicated species and error bars show the standard deviation (n = 2). For each independent experiment, the mean was calculated from duplicate measurements and used in subsequent calculations.

Consistent with sequential transcription, the relative levels of NS1 mRNA decreased for all isolates after four hours pi (GA1: −12%; ON: −5%; GB1: −6%; BA: −22%). Percent reductions were greater for the two RSV isolates, GA1 and BA, with lower total mRNA at four hours pi (avg. of GA1 and BA = 170 ± 70 mRNA/reaction vs. avg. of ON and GB1 = 890 ± 170 mRNA/reaction).

All four sets of steady-state mRNA levels were non-gradient, with levels of G mRNA exceeding levels of N mRNA (Fig 3). Steady-state mRNA levels also showed both subgroup- and genotype-specific differences (Fig 3). Between subgroups, relative levels of NS1 and NS2 were most different (Fig 3), with the two being similar in RSV/A, and with NS1 levels exceeding NS2 by a factor of ∼5 in RSV/B (Fig 3). Within RSV/A, the level of NS1 exceeded NS2 in the GA1 isolate, and was matched by NS2 in the ON isolate (Fig 3). In RSV/B, the level of G mRNA exceeded N in the BA isolate (∼5-fold greater) more than it did in the GB1 isolate (∼2-fold greater) (Fig 3). Furthermore, genotype-specific steady-state mRNA levels were comparable in A549, Vero, and HEp2 cell lines (Fig 4A).

**Fig 3.**
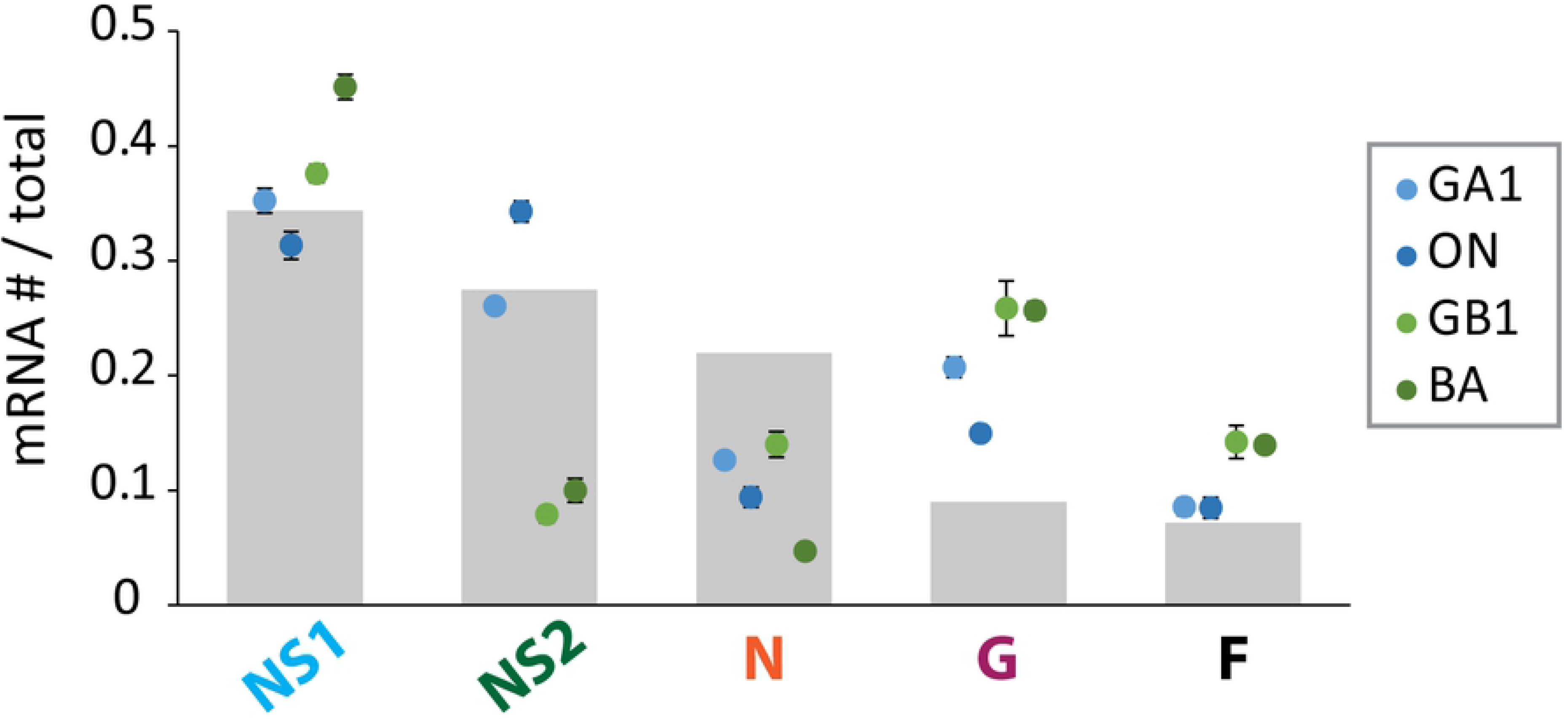
Relative mRNA levels are genotype-specific and non-gradient. Grey bars depict relative mRNA levels expected from an expression gradient resulting from a 20% decrease in transcription at every gene junction. Each dot depicts the mean mRNA # / total mRNA # observed for the indicated species and isolate (RSV/A/GA1Tracy [pale blue]; RSV/A/ON/121301043A [dark blue]; RSV/B/GB1/18537 [light green]; RSV/B/BA/80171 [dark green]) in HEp-2 cells (MOI = 0.01) at steady-state. Steady-state mean relative mRNA levels and standard deviation were calculated using the mean of each relevant time-point (24, 48, 72 hours post-infection). The mean of each time-point was calculated from two independent experiments, and the mean from each experiment was calculated from duplicate measurements as described.

**Fig 4.**
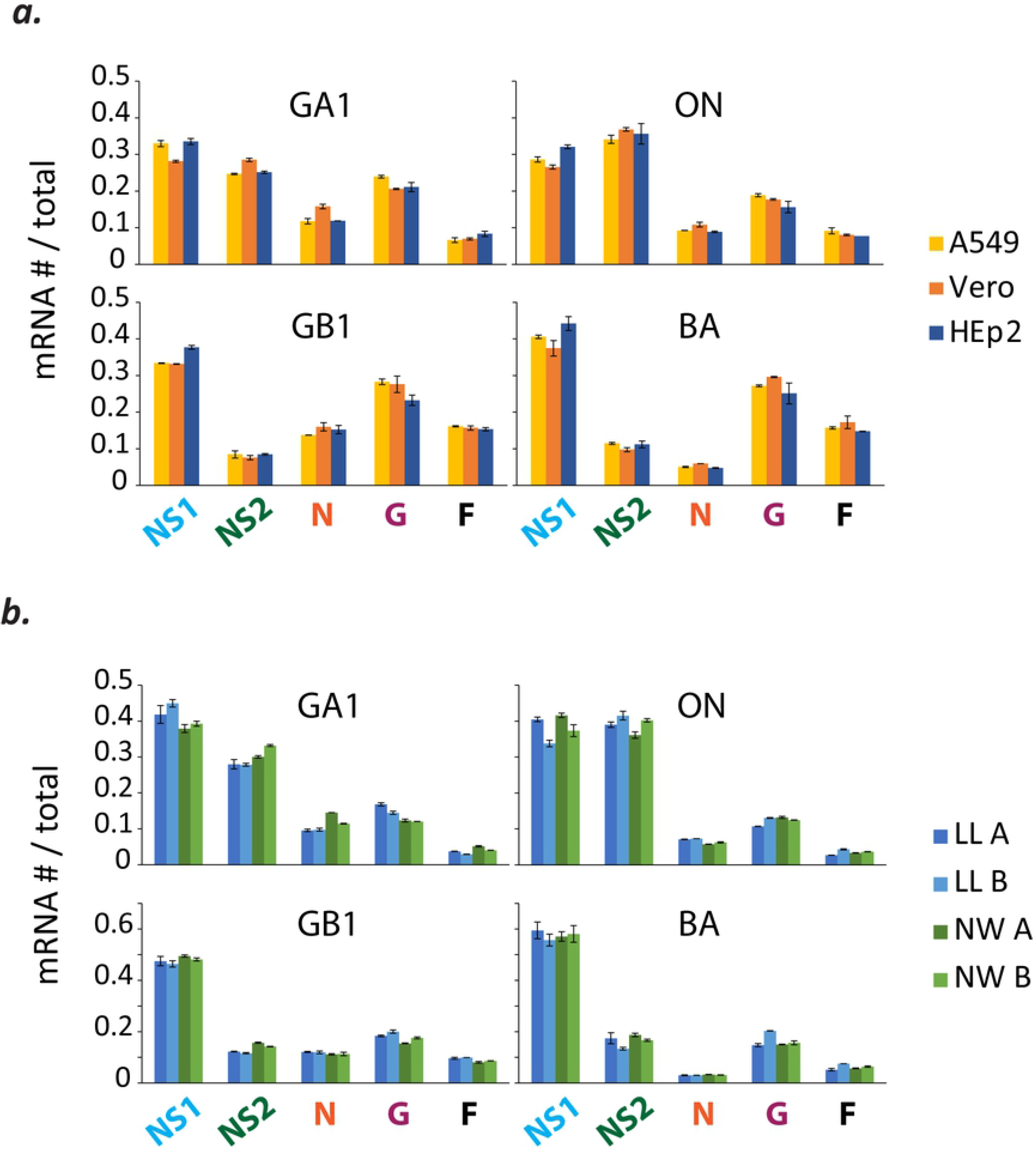
Relative mRNA levels are comparable in different cell lines and in nasal wash and lung lavage samples from infected cotton rats. **(a)** Relative mRNA levels are comparable in different cell lines. Viral mRNA levels were measured from infected A549 (in yellow), Vero (in orange), and HEp-2 (in blue) cell lines (MOI = 0.01) at 24 hours post-infection (pi). Each bar depicts the mean mRNA # / total mRNA # of the indicated species and error bars show the standard deviation (n = 2). For each independent experiment, the mean was calculated from duplicate measurements and used in subsequent calculations. **(b)** Relative mRNA levels are comparable in lung lavage (LL) and nasal wash (NW) samples from infected cotton rats. Each bar depicts the mean mRNA # / total mRNA # of the indicated species and error bars show the standard deviation calculated from duplicate measurements of the same sample. Results from LL samples collected 4 days pi are shown in blue (cotton rat A = light blue; cotton rat B = dark blue) and NW samples shown in green (cotton rat A = light green; cotton rat B = dark green).

We explored whether relative mRNA levels might change in the context of a fully immunocompetent host. A pair of cotton rats was infected with each virus isolate and both lung lavage (LL) and nasal wash (NW) samples were collected at four days pi. Relative mRNA levels were genotype-specific and similar in cotton rat LL and NW samples, and comparable to those measured *in vitro* (Fig 4B).

### RSV mRNA stabilities

The observed divergence from a transcription gradient could be the result of differential stability of the RSV mRNAs. Therefore, we measured transcript stabilities by blocking transcription using the RSV RNA-dependent RNA polymerase (RdRp) inhibitor GS-5734 then monitoring mRNA levels by qPCR over time. Decay was measured for all five mRNAs from each of the four isolates in HEp-2 cells (Fig 5A). Exponential decay functions were fit to the data and half-lives were calculated from the decay constants. Half-lives ranged from 10 to 27 hours with a mean of 16 ± 5 hours (Fig 5B). Distributions of mRNA stabilities varied among the isolates, with GA1 having the greatest uniformity and lowest mean (= 12 ± 1 hours) (Fig 5A). Gene expression patterns were estimated by correcting measured mRNA abundances for degradation and recalculating relative mRNA levels (mRNA expressed = measured mRNA # * *e*^(decay constant * 24 hr)^). Estimated levels of gene expression remained non-gradient; thus, differential mRNA stabilities do not account for the non-gradient patterns observed (Fig 5C). These data indicate that relative mRNA levels are 1) more strongly shaped by gene expression than decay and 2) can safely be interpreted to reflect levels of gene expression.

**Fig 5.**
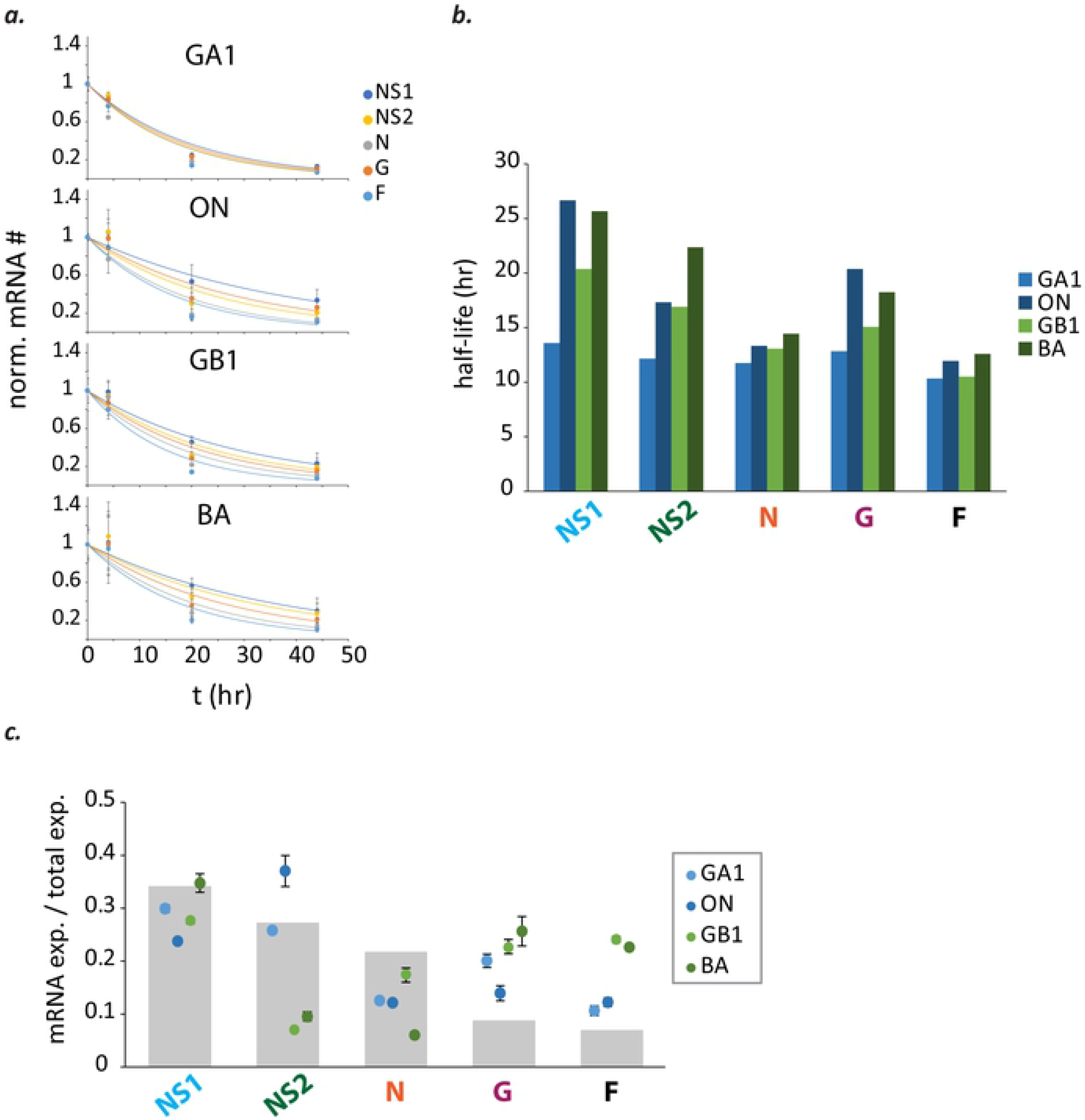
Transcript stabilities do not account for non-gradient patterns, indicating that relative mRNA levels strongly reflect RSV gene expression. **(a)** Viral mRNAs decay after addition of GS-5734, a viral polymerase inhibitor. Viral mRNA levels were divided by RNase P mRNA levels to control for well-to-well variation in the amount of sample obtained, then normalized. Each dot represents the mean normalized mRNA # and error bars the standard deviation of two independent experiments (n=2). For each independent experiment, a mean was calculated from the means of two different samples; and each sample mean was obtained from duplicate measurements. **(b)** Decay constants obtained from exponential decay functions fit to each data set were used to calculate mRNA half-lives (RSV/A/GA1Tracy [pale blue]; RSV/A/ON/121301043A [dark blue]; RSV/B/GB1/18537 [light green]; RSV/B/BA/80171 [dark green]). **(c)** Transcript stabilities cannot account for non-gradient mRNA levels. Grey bars depict relative mRNA levels expected from an expression gradient resulting from a 20% decrease in transcription at every gene junction. Each dot depicts the mean expressed mRNA # / total expressed mRNA # estimated for the indicated mRNA species and virus isolate in HEp-2 cells (MOI = 0.01) at 24 hours post-infection (mRNA expressed = mRNA # observed * *e*^(decay constant * 24 hr)^).

### GS signal sequence and N-phase

Whole genome sequences of the four RSV isolates were obtained by next-generation sequencing and analyzed for differences in GS signals that might help explain the non-gradient gene expression patterns observed. GS signals were highly conserved, with a single U to C substitution at position ten of the G gene GS signal (Fig 6A).

**Fig 6.**
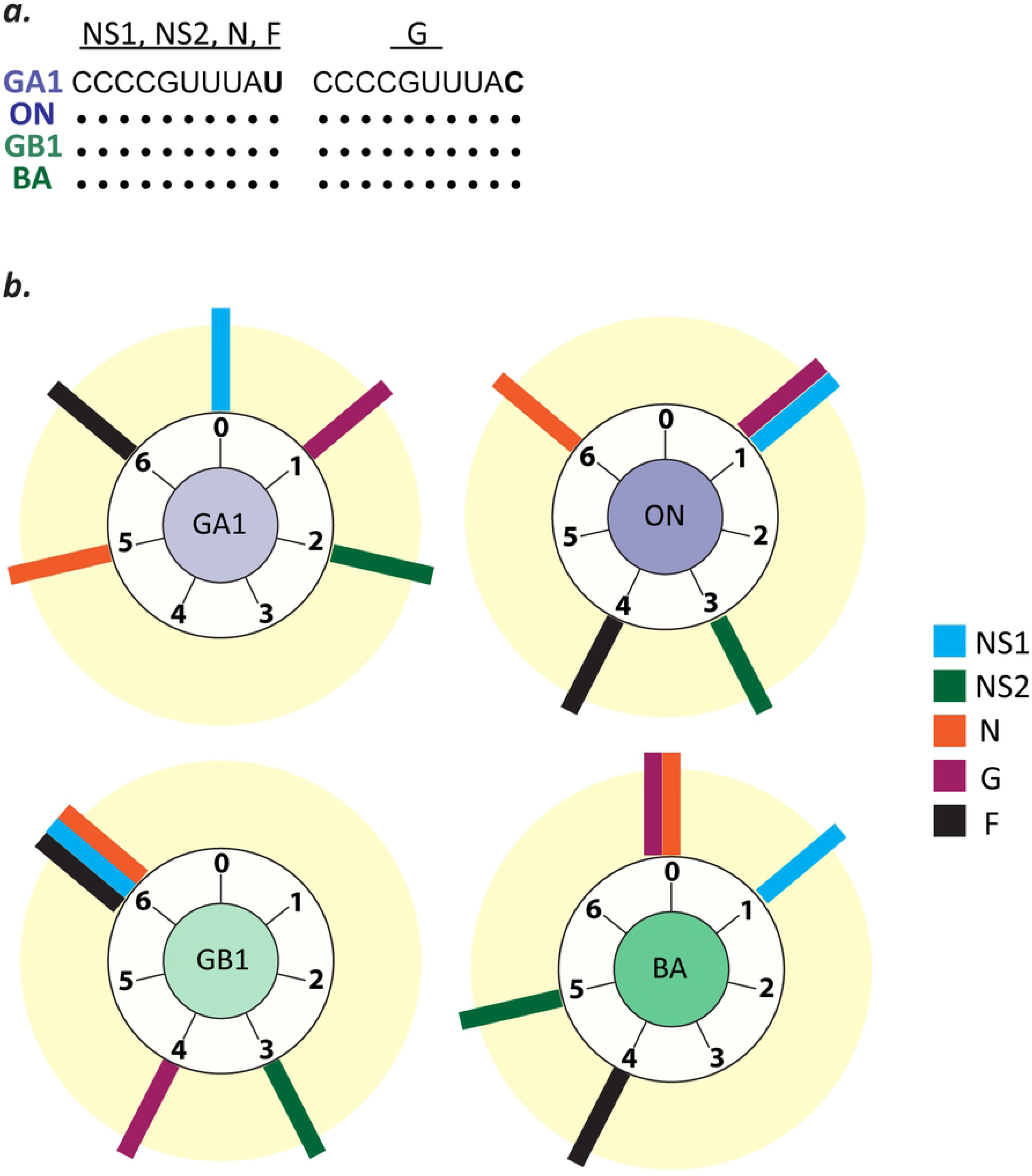
GS signal sequence is highly conserved but alignment with N protein (N-phase) is variable, revealing a potential source of genotype-specific and non-gradient gene expression patterns. **(a)** GS signals are highly conserved and show only a U to C substitution at position ten of the G gene GS signal. (Genomic, i.e., negative-strand, sequence displayed). **(b)** GS signals have variable N-phase. Diagrams show the estimated GS signal N-phase for each gene whose mRNA levels were measured (NS1 (cyan), NS2 (green), N (tawny), G (purple), & F (charcoal)) from the 4 virus isolates used.

We analyzed GS signal sequences for their alignment with N protein, as the alignment of a GS signal with bound N protein will affect its conformation and determine its configuration of solvent-exposed and buried nucleobases [16]. The alignment of a GS signal with N protein (N-phase) might therefore affect interactions with scanning polymerases and alter the likelihood of transcription initiation. We estimated the N-phase of each GS signal by calculating the remainder resulting from dividing the number of nucleotides separating the GS signal from the L GE signal by seven (the number of nucleotides bound by one subunit of N). The L GE signal was used as a proxy as the exact 5’ terminus of each RSV genotype is not known. Thus, the estimated GS signal N-phase will differ from the actual N-phase if the nucleotide length beyond the end of the L GE signal is not equal to an integer multiple of seven. However, every GS signal N-phase within a genotype would be uniformly affected, making estimated intra-genotype differences equal to actual intra-genotype differences. GS signal N-phase was highly variable, making it a potential source of the variation observed in patterns of gene expression (Fig 6B).

### Minigenomes to assess the effect of changing GS signal N-phase on gene expression

We hypothesized that changing GS signal N-phase would alter the likelihood of transcription initiation. To test our hypothesis, we designed plasmids encoding RSV minigenomes containing two reporter genes (Renilla luciferase and Firefly luciferase) each flanked by GS and GE signals [45, 46] (Fig 7A). We specifically altered the N-phase of the Firefly GS signal by introducing single nucleotide insertions within the adjoining 5’ untranslated region (UTR) along with compensatory single nucleotide deletions within the adjoining intergenic region (Fig 7B). In this way, both the total length of the minigenome and the N-phase of all other sequences were fixed (Fig 7B). If gene expression occurred independently of GS signal N-phase, ratios of luciferase activity would remain constant. Measured ratios of luciferase activity showed a switch-like dependence on GS signal N-phase, with four states resulting in relatively high activity, two states with low, and one state with intermediate activity (Fig 7C). Ratios increased by as much as 50% relative to the minimum measured (Fig 7C). Thus the N-phase of the Firefly luciferase GS signal affected the relative level of gene expression, and by inference, transcription initiation (Fig 7C). Furthermore, ratios of luciferase activity were consistent with a periodicity of seven nucleotides (Fig 7C).

**Fig 7.**
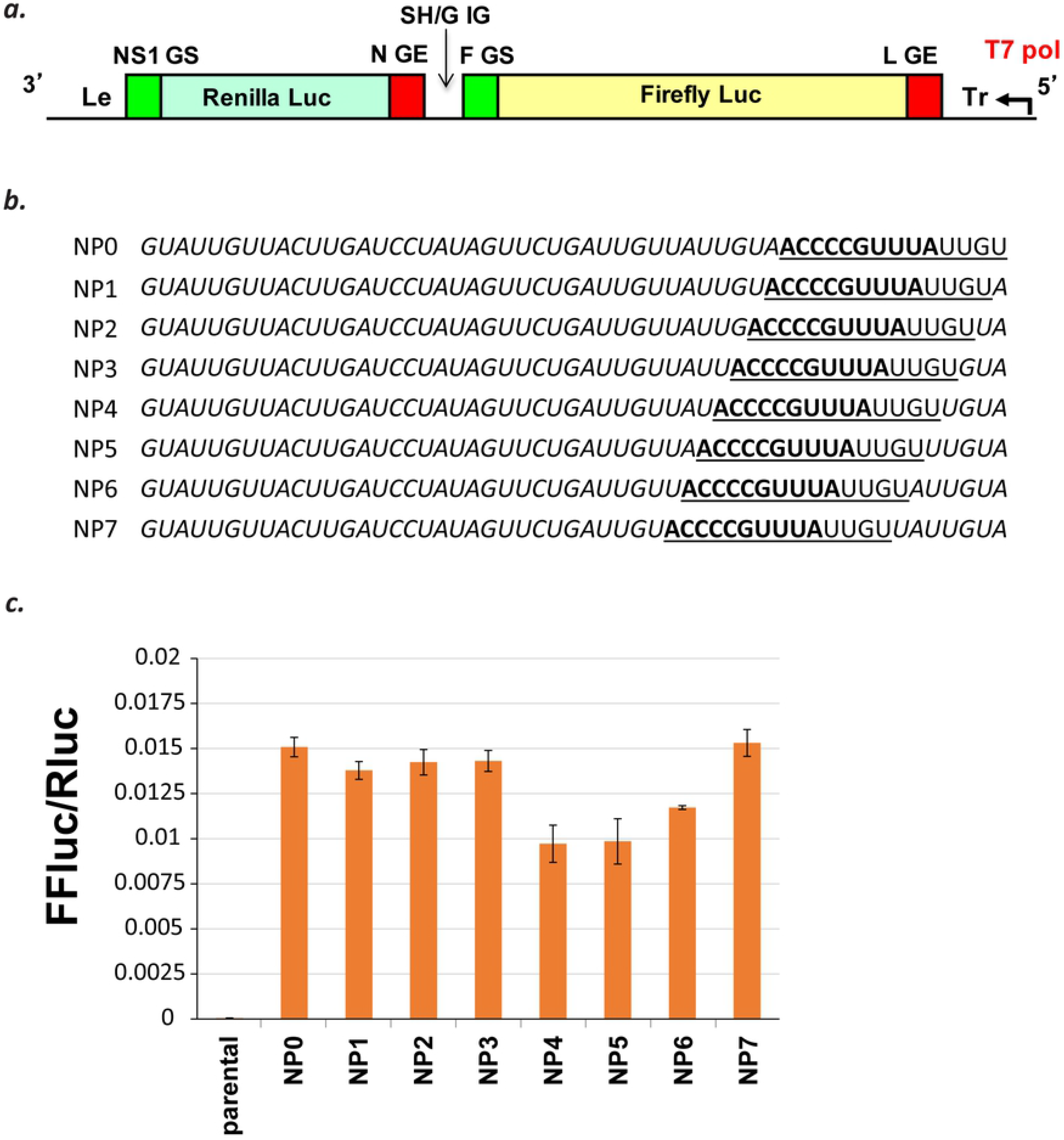
Gene expression changes with GS signal N-phase in minigenomes encoding two luciferase enzymes. **(a)** A minigenome encoding two reporter genes, the second with variable GS signal N-phase. RSV minigenomes contained Renilla and Firefly luciferase genes separated by an N GE signal, SH/G intergenic (IG) sequence, and F GS signal of variable N-phase. **(b)** The F GS signal and neighboring sequences in minigenomes with variable GS signal N-phase. In *italics*: SH-G intergenic (IG) sequence; in **bold**: consensus GS signal sequence; <u>underlined</u>: F GS signal sequence block. (Genomic, i.e., negative-strand, sequence displayed). **(c)** Changing ratios of Firefly to Renilla luciferase activity indicate that GS signal N-phase can affect gene expression. Each bar represents the mean and error bars the standard deviation of measurements of four samples taken at 24 hr pi from a single experiment.

## Discussion

We observed genotype-dependent and non-gradient patterns of RSV gene expression. We hypothesize that non-gradient patterns require a mechanism to alter the likelihood of transcription initiation at different GS signals. GS signal sequences were highly conserved but varied in alignment with N protein, or N-phase, providing a potential source of biased transcription initiation. Using RSV minigenomes, we showed that varying GS signal N-phase can affect gene expression. These unexpected findings highlight gaps in our knowledge of RSV transcription and raise important issues relevant to future studies.

Accurate mRNA abundance measurements by qPCR require reagents that bind target without any mismatches [47, 48]. Perfectly designed and distinct sets of reagents can amplify target with variable efficiency, as the amplification efficiency depends on the physicochemical properties of the reagents (the free energies of different intra- and intermolecular interactions) and the qPCR conditions used. For our 20 oligonucleotide standards, we found the lowest melting temperature from each set of reagents correlated positively with amplification efficiencies and negatively with cycle threshold values (S1 Fig). These correlations indicate that physicochemical differences in the primers and probes can account for the minor variation observed in the amplification of oligonucleotide standards, and support the accuracy of our approach to measuring viral mRNA abundances.

A gene expression gradient has been widely assumed for RSV, but supporting data come from a modest number of studies and are largely restricted to laboratory-adapted isolates (Long and A2) from the prototypic GA1 genotype of subgroup A. The first measurements were made by Collins and Wertz (1983) using an A2 strain in HEp-2 cells [28, 42, 49]. They discovered the gene order of RSV and found it was approximated by decreasing mRNA abundances measured by northern blot [28, 42, 49]. Barik later reported a gradient by dot blot hybridization of radiolabeled mRNAs produced *in vitro* using ribonucleoprotein (RNP) complex from an RSV Long strain and cell extract from uninfected HEp-2 cells [41]. Over a decade later, Boukhvalova et al. measured a gradient-like pattern by qPCR of mRNA abundances from an RSV Long strain grown in A549 cells [39]. In contrast, Aljabr et al. recently reported mRNA abundances by RNA-Seq from an A2 strain in HEp-2 cells that are inconsistent with a gradient. The most abundant mRNA they observed was associated with the G gene [40]. Levitz et al. reported the G gene to be the most highly expressed gene at later time-points in A549 cells infected with isolates from the RSV/B subgroup [43]. Thus, recent published data indicate that patterns of RSV gene expression vary and do not always follow a gradient. Here, we report data from isolates belonging to four different genotypes (GA1, ON, GB1, BA) showing variable and non-gradient patterns of gene expression and all with an apparent excess of G mRNA.

Studies of transcription in RSV and other NNS RNA viruses show that gene expression depends on a variety of factors. Among the factors affecting RSV gene expression are sequences of GS and GE signals [30–32], intergenic (IG) sequences that can change how efficiently transcription is terminated or initiated [50, 51], and other factors including polymerase mutations and sequences of unknown function [52, 53]. Our minigenome experiments add GS signal N-phase to the list of factors involved in RSV gene expression.

Our minigenome data suggest that polymerases preferentially initiate transcription at GS signals with certain solvent-exposed nucleobases (3C and 10U of the RSV GS signal). What accounts for this preference, and what events follow GS signal recognition and lead to either transcription initiation or continued scanning is unknown. It is interesting that the U to C substitution in position ten of the G gene GS signal has been shown to result in less not more transcription [30]. Thus, additional factor(s) beyond GS signal N-phase may account for over-expression of the G gene. It is worth stating that transcription initiation, being a molecular event, must be stochastic. RSV transcription is therefore sequential but likely *not obligatorily* sequential. A relative excess of G gene mRNA can occur from polymerases, more often than not, failing to initiate transcription at the N gene before initiating at the G gene. It is also possible that the N gene is usually expressed before the G gene, but G mRNA accumulates more because of polymerase scanning and increased re-initiation of transcription at G. Either scenario might contribute and both are consistent with the ultraviolet (UV) transcriptional mapping data underlying sequential transcription [24–26, 28].

Differences in luciferase activities from minigenomes are smaller than the differences observed in viral mRNA levels. Several factors might help explain this finding. First, the viral polymerase complex proteins (M2-1, N, P, L) that drive RNA synthesis in the minigenome system are over-expressed by transfection of plasmids encoding codon-optimized genes. This over-abundance of polymerase proteins might not accurately represent the situation during RSV infection. Minigenomes are also proxies for full-length genomes, being shorter and having a simpler genetic structure. For instance, dicistronic minigenomes contain one pair of GS and GE signals straddling a single intergenic sequence while genomes contain nine variably spaced pairs of GS and GE signals straddling nine intergenic sequences of variable sequence and length. Interactions of transcribing and scanning polymerases with the structurally complex RSV genome might, in nonobvious ways, bias the expression of some genes over others. There could also be differences in the stability of the mRNAs or proteins for Renilla versus Firefly luciferase. Finally, concentrations of both luciferase proteins may have reached saturating intracellular levels prior to measurement (24 hr pi). Saturating protein concentrations would underrepresent differences in mRNA levels.

Our results show that transcription initiation by the RSV polymerase depends in part on GS signal N-phase. This potentially helps explain our and other recent observations of 1) non-gradient and 2) variable patterns of gene expression. The functional importance of genotype-dependent patterns of gene expression demands exploration. Finally, GS signal N-phase-regulated transcription initiation might also play a role in other NNS viruses.

## Materials and Methods

### Virus strains

RSV isolates were initially genotyped as described [13, 54] by sequencing a 270 bp fragment in the second hypervariable region of the G gene. RSV/A/GA1/Tracy and RSV/B/GB1/18537 are prototypic strains isolated in 1989 and 1962, respectively [13], while RSV/A/ON/121301043A and RSV/B/BA/80171 are contemporaneous strains isolated in 2013 and 2010, respectively [55, 56].

### Cell-lines and cotton rats

HEp-2 (ATCC CCL-23), A549 (ATCC CCL-185), and Vero (ATCC CCL-81) were cultured in minimal essential medium (MEM) containing 10% fetal bovine serum (FBS), 1 μg/ml penicillin, streptomycin, and amphotericin B (PSA), and supplemented with L-glutamine.

Male and female *Sigmodon hispidus* cotton rats were bred and housed in the vivarium in Baylor College of Medicine. Cotton rats were ∼75 to 150 g of body weight at the start of the experiments.

### Viral replication in cell culture and cotton rats

The media from 70-90% confluent HEp-2, A549, or Vero cells in 24-well plates was aspirated, and 0.2 ml of virus diluted in MEM containing 2% FBS with antibiotics, antifungal, and L-glutamine (2% FBS-MEM) was added to replicate wells for each of the time-points to be acquired. Plates were incubated at 37°C and 5% CO_2_ for 1 hour. Following infection, virus-containing media was aspirated and replaced with 1 ml of pre-warmed 2% FBS-MEM. Plates were incubated at 37°C and 5% CO_2_ until sample collection. At each time point, the media was aspirated and infected monolayers were lysed with 1X RIPA buffer and pelleted by centrifugation. The supernatant was flash frozen in a mixture of dry ice and 95% ethanol then stored at −80°C.

Eight- to ten-week-old male and female cotton rats were sedated and inoculated intranasally with 10^5^ plaque forming units (pfu) of RSV as described [57]. Cotton rats were euthanized on day 4 post-infection. Nasal wash (NW) samples were collected from each cotton rat by disarticulating the jaw and washing with 2 ml of collection media (= Iscove’s media containing 15% glycerin and mixed 1:1 with 2% FBS-MEM) through each nare, collecting the wash from the posterior opening of the pallet. Lung lavage (LL) samples were collected after the left lung lobe was removed and rinsed in sterile water to remove external blood contamination and weighed. The left lobe was transpleurally lavaged using 3 mL of collection media. Both NW and LL fluids were stored at −80°C.

### RNA extraction and reverse transcription

Viral RNA was extracted from clarified cell lysates or samples obtained from cotton rats as described [55] by using the Mini Viral RNA Kit (Qiagen Sciences, Germantown, Maryland) and automated platform QIAcube (Qiagen, Hilden, Germany) according to the manufacturer’s instructions. Complementary DNA (cDNA) was generated using the SuperScript™ IV First-Strand Synthesis System and oligo(dT)_20_ primers according to the manufacturer’s instructions (ThermoFisher Scientific).

### RSV mRNA abundance measurements

Accurate mRNA abundance measurements by qPCR require reagents that bind target without any mismatches [47, 48]. Twenty sets of target-specific primers and probes (from five mRNA targets for four virus isolates) were designed using whole genome sequences obtained by next-generation sequencing. *C*_T_ values were measured using the StepOnePlus Real-Time PCR System (ThermoFisher Scientific). Thresholding was performed according to the manufacturer’s instructions [58].

Oligonucleotide standards were used to convert sample *C*_T_ values to mRNA abundances. Twenty oligonucleotide standards identical in sequence to the 20 targets of the specific primers and probes described above were purchased from IDT^®^, received lyophilized and resuspended in TE buffer pH 8. Each oligonucleotide standard was diluted to 4×10^6^ molecules/μl and further diluted serially to a concentration of 40 molecules/μl in TE buffer. Duplicate *C*_T_ values were measured for each dilution and an average *C*_T_ was calculated. Average *C*_T_ values and known amounts (molecules/rxn) were used to construct a standard curve for each oligonucleotide standard.

For cDNAs derived from *in vitro* cell lysate or cotton rat samples, *C*_T_ values were measured in duplicate and used to calculate an average. Each average sample *C*_T_ value was converted to an mRNA abundance using the linear relationship determined for the appropriate oligonucleotide standard *C*_T_ vs. log10 of the oligonucleotide standard amount (molecules/rxn).

### RSV mRNA stability measurements

Samples of HEp-2 cells infected with virus isolates at an MOI of 0.01 were collected from single wells of 24-well plates at multiple time-points up to 48 hours after addition of 100 μM GS-5734. GS-5734 is a monophosphate prodrug of an adenosine nucleoside analog that binds a broad range of viral RNA-dependent RNA polymerases (RdRps) and acts as an RNA chain terminator [59, 60]. Samples were collected as described above using 1X RIPA buffer to lyse infected cells, clarifying the lysate by centrifugation, and flash-freezing and storing the clarified lysate at −80°C. Viral RNA were extracted and converted to cDNA using oligo(dT)_20_ primers. Transcript levels from RNase P (a host housekeeping gene) were measured using qPCR reagents acquired from the Centers for Disease Control and Prevention (CDC) and used to correct viral mRNA levels for well-to-well variation in the amount of sample obtained. Exponential decay functions were fit to the normalized data and used to calculate half-lives. Estimates of the amounts of mRNA expressed up to 24 hours pi were made by correcting the observed mRNA abundances at 24 hours pi for degradation using the exponential decay constants calculated (the number of expressed = the number of observed * *e*^(decay constant * 24 hr)^) and assuming production of all observed mRNA at t = 0 hours post-infection. This unrealistic assumption maximizes the effect of different rates of decay on the estimated levels of total expressed mRNA.

### Whole genome sequencing and assembly

cDNAs for sequencing were generated from viral RNA using the SuperScript™ VILO™ cDNA Synthesis Kit and random hexamers (ThermoFisher Scientific). cDNAs were amplified using specific primers, and PCR products of each sample were purified and pooled [61]. Pooled PCR products (1 μg) were digested with the NEBNext dsDNA fragmentase kit (New England BioLabs, Inc., Ipswich, MA). Fragmented DNA was end-repaired with the NEBNext End Repair Module (New England BioLabs, Inc.). End-repaired DNA was ligated with the Ion P1 adaptor and unique Ion Xpress barcode adaptors (KAPA Adapter Kit 1–24; KAPABiosystems). Agencourt AMPure XP beads (Beckman Coulter, Inc., Brea, CA) were used to selectively capture DNA between 100 and 250 bp in length. All reaction products were purified with the Isolate II PCR kit (Bioline USA, Inc.). These libraries underwent nick translation and amplification. Experion Automated Electrophoresis System (Bio-Rad Laboratories, Inc., Hercules, CA) was used to confirm fragment lengths and molar concentrations. Equal molar amounts of all libraries were pooled and libraries were sequenced by Ion Proton™ System (ThermoFisher Scientific) generating 150 bp reads. Raw data, FASTQ and BAM files, were generated by the Torrent Suite™ Software (version 5.0.4; ThermoFisher Scientific).

Reads were assembled by Iterative Refinement Meta-Assembler (IRMA), which was designed for highly variable RNA viruses with more robust assembly and variant calling [62, 63]. IRMA v0.6.7 (https://wonder.cdc.gov/amd/flu/irma/) was used with an assembly module specifically designed for RSV.

### Estimates of gene start (GS) signal N-phase

GS signal N-phase was estimated from whole genome sequences using the end of the L (polymerase) GE signal [30]. The L GE signal (3’-UCAAUAAUUUUU-5’; genome sense) was used as a proxy for the 5’ end of the RSV genome. N-phase was calculated by determining the number of bases between the last U of the L GE and the first C of the GS (3’-CCCCGUUUAU-5’) then dividing by seven, representing the number of bases encapsidated by a single N molecule.

### Minigenome experiments

RSV minigenomes contained Renilla luciferase and Firefly luciferase genes flanked by the RSV A2 leader (le)-NS1 GS and L GE-trailer (Tr) sequences [45, 46]. The Renilla and Firefly luciferase genes were separated by an N/M GE signal, SH-G intergenic (IG) sequence, and F GS signal of variable N-phase. The N-phase of the F GS signal was altered by sequentially deleting the nucleotide immediately 3’ to the GS and inserting the same nucleotide immediately 5’ of the F GS signal. Eight constructs were constructed assuming the periodicity of seven nucleotides established by Tawar et al. [16]. Minigenomes were co-transfected into BSR-T7 cells with expression plasmids encoding codon-optimized N, P, L, and M2-1 genes [45, 46]. Firefly and Renilla luciferase activities were measured at 24 hours post-transfection using Dual Luciferase Reagent (Promega) [46].

### Ethics Statement

All experimental protocols were approved by the Baylor College of Medicine’s Institutional Animal Care and Use Committee (IACUC) (license # AN-2307). All experiments were conducted in accordance with the Guide for Care and Use of Laboratory Animals of the National Institutes of Health, as well as local, state and federal laws.

### Accession numbers

Sequences reported in this study were deposited in GenBank database under accession numbers MG813977-MG813995.

## Acknowledgements

Thanks to Michel Perron of Gilead for providing the viral RdRp inhibitor GS-5734 for use in experiments to measure RSV transcript stabilities. Thanks to Brian Gilbert from the Department of Molecular Virology and Microbiology at Baylor College of Medicine for providing the cotton rats needed to perform RSV infection studies. We thank Kim Tran for technical assistance.

## Supporting Information Legends

**S1 Fig. Amplification efficiencies positively correlate and *C*_T_ values negatively correlate with the minimum melting temperature (min. T_m_) of the target-specific qPCR reagents used. (a)** Pearson correlation for amplification efficiencies vs. min. Tm: *R*=0.57, *p*=0.0086. **(b)** Pearson correlations for *C*_T_ values measured at the extremes of target quantity (200 and 2×10^7^ molecules / rxn) vs. min. Tm: *R*=−0.65, *p*=0.002 and *R*=−0.66, *p*=0.0015, respectively.

